# Genetic and genomic analysis of early abortions in Israeli dairy cattle

**DOI:** 10.1101/557306

**Authors:** Moran Gershoni, Ephraim Ezra, Joel Ira Weller

## Abstract

Female infertility accounts for at least 50% of all human infertility cases. One of the causes contributing for female infertility is embryo loss after fertilization. Previous findings suggested that more than half of fertilizations results in embryo loss before pregnancy is detected. Dairy cattle may be a useful model for study of the genetic architecture of this trait. In advanced commercial populations, all breeding is by artificial insemination, and extensive records of the cows’ estrus, insemination and pregnancies are available. We proposed re-insemination between 49 and 100 days after the first insemination as an indicator trait for early abortion in dairy cattle, based on the mean estrus interval of 21 days. Israeli Holstein cows scored as early abortion were compare to cows recorded as pregnant from the first insemination. This trait was compare to conception rate from first insemination. Animal model variance components were estimated by REML, including parents and grandparents of cows with records. First parity heritability for conception rate was 3%. In the multi-trait analysis of parities 1-3 for abortion rate heritabilities ranged from 8.9% for first parity to 10.4% for second parity. The variance component for the service sire effect for abortion rate were less than half the variance component for conception rate. Thus genetic control of the two traits is clearly different. Genome wide association study were performed based on the genetic evaluations of ∼1200 sires with reliabilities >50%. The markers with the lowest probabilities for early abortion were also included among the markers with the lowest probabilities for conception rate, but not vice versa. The marker explaining the most variance for abortion rate is located within the ABCA9 gene, which is found within an ABC genes cluster. The ATP-binding cassette family is the major class of primary active transporters in the placenta.

**Author summary:** Approximately 70% of human conceptions fail to achieve viability. Almost 50% of all pregnancies end in miscarriage before the clinical recognition of a missed period. Cattle are a useful model for human female reproductive processes, because of the similarities in the reproductive cycles, and the extensive documentation in commercial cattle populations, including estrus and insemination records. In addition to the expected benefits from cow fertility research for human biomedical applications, fertility is an economically important trait in dairy cattle with very low heritability. The mean estrous interval for cattle is 21 days. We therefore proposed re-insemination between 49 and 100 days after the first insemination as an indicator trait for early abortion. Israeli Holstein cows scored as having early abortion based on first insemination after parturition were compare to cows recorded as pregnant from the first insemination. Heritability for early abortion rate was three-fold the heritability for conception rate. In a genome wide association study based on 1200 dairy bulls genotyped for 41,000 markers, six markers were found with nominal probabilities of < 10^-12^ to reject the null hypothesis of no effect on early abortion rate. Early abortion rate may be a useful indicator trait for improvement of fertility in dairy cattle.

## Introduction

Genetic factors that reduce the ability of an individual to reproduce are expected to be under intensive negative selection, and therefore to remain rare in the population. It is thus surprising that in humans about 15% of the reproductive age couples cannot achieve successful pregnancy without medical assistance [1, 2]. This is partially due to the polygenic and sexually differentiate nature of this trait [3, 4]. Female infertility apparently accounts for more than half of the cases [5]. Previous studies have suggested that the chances of a woman to achieve a successful pregnancy per menstrual cycle is approximately 25% [6, 7]. Among other causes, the lack of detected pregnancy could be the result of early abortion (EA). Approximately 70% of human conceptions fail to achieve viability, with almost 50% of all pregnancies ending in miscarriage before the clinical recognition of a missed period, or the presence of embryonal heart activity [8, 9]. This complicates the study of EA in humans, although it is likely to have a large impact on human fertility.

Cattle were found to be a useful model for human female reproductive processes, mainly because of the similarities in the reproductive cycles [10]. Within this context, cows were used in studies on ovarian function [11], effects of ageing on fertility [12], embryo-maternal communication [13, 14], pregnancy maintenance and irregularities associated with assisted reproduction techniques [15, 16]. The extensive documentation in many commercial cattle populations, including estrus and insemination records, provides a good opportunity to investigate the genetics of EA. In addition, each ejaculation is evaluated by AI labs, and defective semen is rejected. Thus, the service sire has only a very minor effect on conception rate [17]. Furthermore, fertility is an economically important trait in dairy cattle [18].

Like all economic traits in dairy cattle, genetic evaluation of fertility is based on field records. Unlike milk production traits and somatic cell concentration, there is no accepted consensus on how fertility should be scored. Traditionally the most common criterion was “non-return rate,” the fraction of cows that were not re-inseminated within a specific time interval. This criterion ignored cows that were culled after the first insemination [19]. More recent measures have considered the time laps from first insemination to pregnancy or some function of the number of times a cow is inseminated during the lactation [20]. Nearly all measures of female fertility have very low heritability, in the range of 1 to 4% [18, 20].

Although abortions are recorded in Israel, very few first trimester abortions are noted either by the herd manager or the attending veterinarian. Indeed, previous studies in other populations estimated fertilization rate as greater than 75%, while the conception rate (CR) was approximately 35% [21]. The differences between these two observations are likely due to pregnancy loss that occur in more than 50%

Since genotyping of large numbers of animals with high density SNP chips has become routine, a number of recessive lethal alleles have been detected in commercial dairy cattle populations, which result in early term abortions [25]. Detection was originally based on the lack of homozygotes for the haplotype harboring the lethal allele, and a reduction in fertility rate for daughters of sires that received this haplotype. In several cases the causative polymorphism has been determined, e. g. [26]. Since the abortion is generally not observed, pregnancies of fetuses homozygous for the lethal alleles are recorded as “non-conception.”

Many studies have shown that the average estrous interval in dairy cattle is 21 days [27]. Thus, most cows that do not conceive in the first service should be re-inseminated approximately 21 days later. The number of days between first and second insemination for first parity were previously shown with a peak at 21 days, and a secondary peak around 42 days, which corresponds to 2 estrus cycles. Despite these two prominent peaks, a significant number of cows were re-inseminated at >45 days after the first insemination [28]. Although these late inseminations may be due to non-conception at the first insemination, and lack of observed estrous at the expected interval; another explanation is: conception at the first insemination, and early term abortion due to embryonic lethality or female factors increasing the predisposition for embryonic death.

One way to test this hypothesis is to demonstrate that the genetic factors that control long intervals between first and second inseminations are different from the genetic factors that control conception, and to evaluate the genetic contributions of the cow and the service sire. The objectives of this study were a genetic and genomic analysis of cows with long intervals between first and second insemination, as compared to cows recorded as conceiving at first insemination; and comparison of this trait to CR at first insemination.

## Results

To define long interval between inseminations as indication for EA, we first analyzed the distribution of the insemination interval in Israeli-Holstein cows that were inseminated more than once (Fig 1). The distribution of the insemination interval was similar to previously reported [28]. Therefore, we defined EA as occurring when the cow was re-inseminated between 49-100 days after the first insemination (see also in the material and methods section). This interval was previously suggested to represent embryonic death in most instances [29].

**Fig 1.**
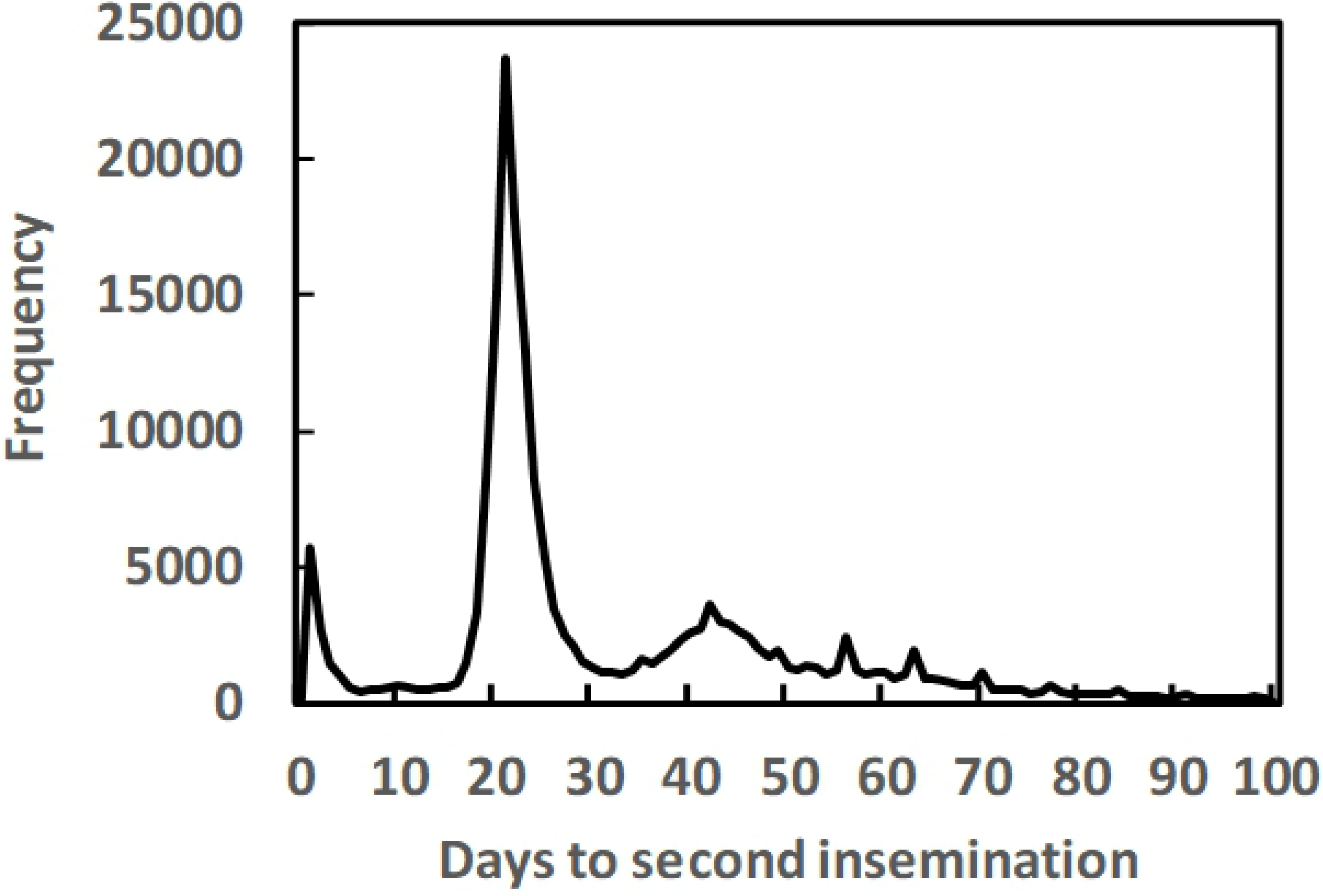
Frequencies of days between first and second inseminations for first parity Israeli Holstein cows. Valid records of cows freshening between 2007 and 2016 are included.

Effects of insemination month on conception and abortion rate as computed from the REML analyses of data sets one and two are shown in Fig 2. Effects were set to zero for December. Generally, the effects were similar for both traits, with major reductions in the late summer, August and September. These results correspond to previous results for the effect of insemination month on CR of Israeli Holsteins [17].

**Fig 2.**
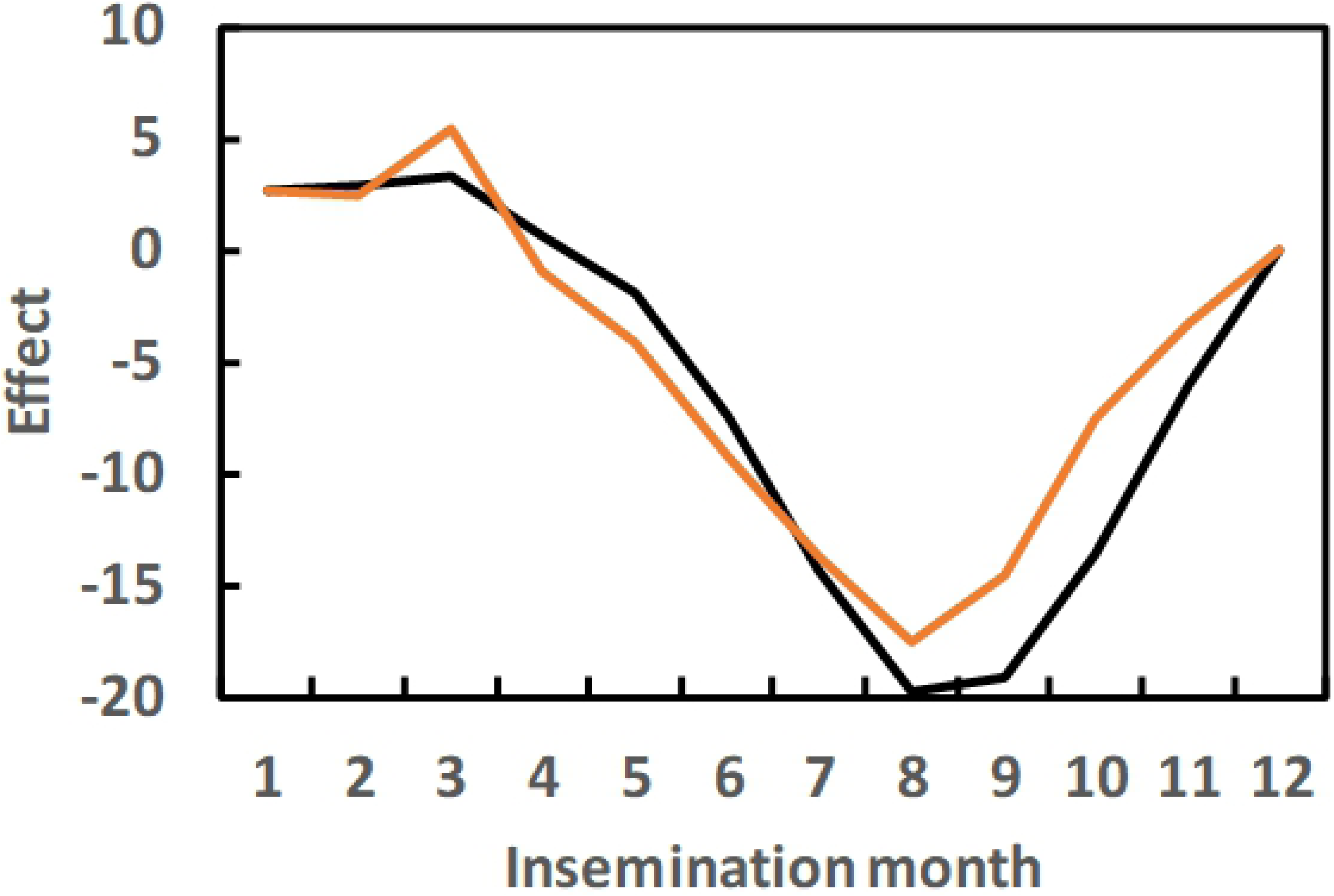
Effects of insemination month on conception and abortion rate. Based on valid records of cows freshening between 2007 and 2016. 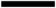, Conception rate; 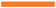, early abortion rate. A score of 0 = non-conception or abortion, and 100 = normal pregnancy. Effects are computed relative to December.

Estimates of variance components from the REML analyses of data sets one through four, and the heritabilities are given in Table 3. First parity heritabilities were 3.0% for CR from first insemination, but 7.7% for EA. For CR with assumed abortions deleted from the analysis, heritability for CR decreased to 2.6%. In the multi-trait analysis of parities 1-3 for EA heritabilities ranges from 8.9% for first parity to 10.4% for second parity, but differences among the parities were not significant. The variance components for the service sire were 8.6 and 10.1 for the two analyses of conception, but ≤ 3.5 for all the EA analyses. Although the service sire can effect EA by transfer of recessive lethal alleles to the fetus [25], the service sire factor apparently does not explain a major proportion of the genetic variance for EA. The additive genetic variance of the inseminated cow for EA is ∼50 times greater than the variance of the service sire, as opposed to 8-fold for CR. Thus genetic control for the two traits is clearly different.

**Table 3.**
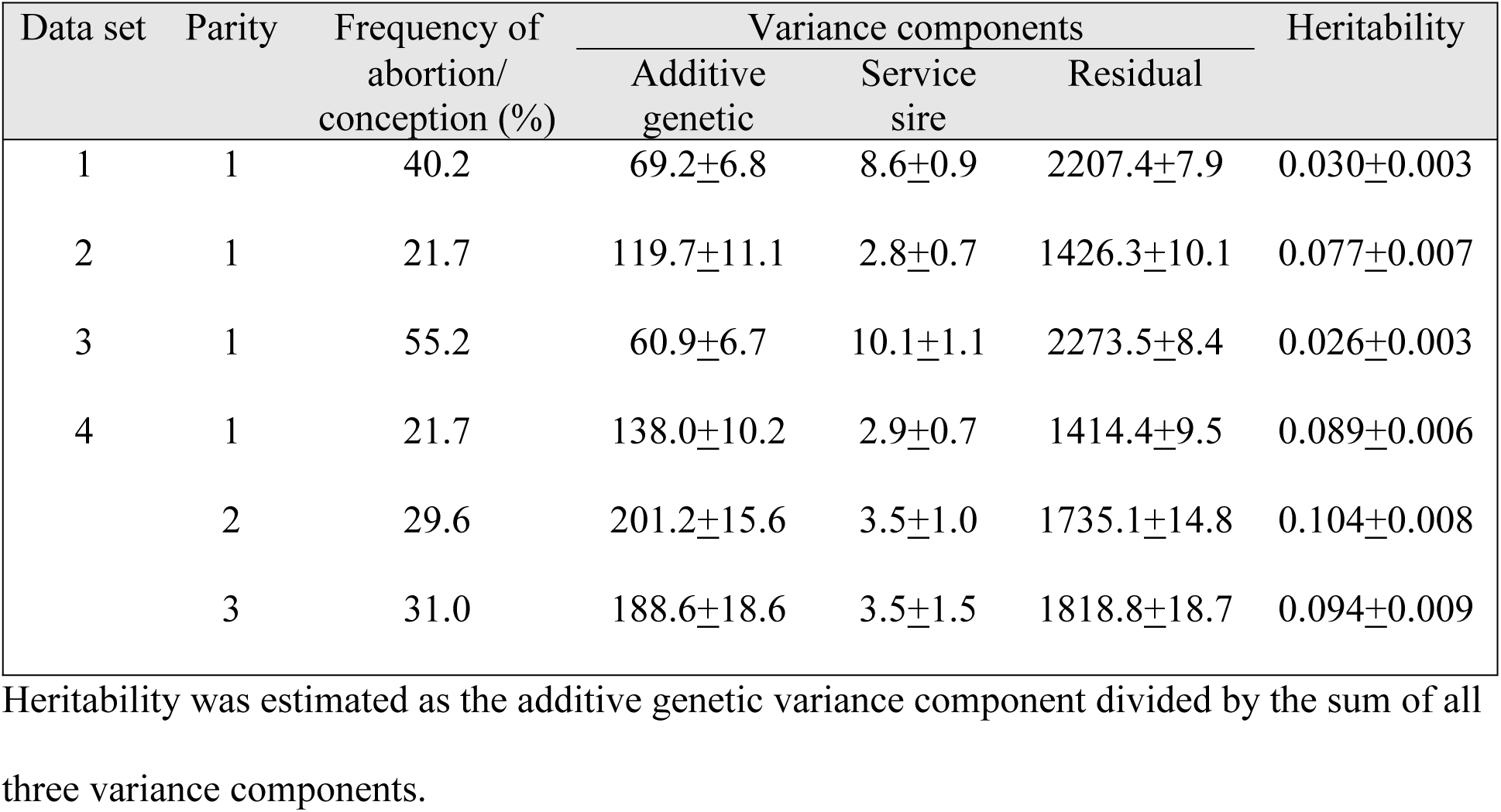
Variance components (± standard errors) and heritabilities for data sets 1-4. Heritability was estimated as the additive genetic variance component divided by the sum of all three variance components.

The genetic and environmental correlations among the three parities for the data set 4 analysis are given in Table 4. All genetic correlations were > 0.9, while all environmental correlations were < 0.12. Thus analysis by the single trait animal model is justified, with the variance ratios given in the methods section.

**Table 4.** Genetic and environmental correlations ± standard errors for abortion rate (data set 4). Genetic correlations were the correlations among the additive genetic effects, and the environmental correlations were the correlations among the residual effects.

| Parities | Correlations |  |
| --- | --- | --- |
|  | Genetic | Environmental |
| 1, 2 | 0.966±0.012 | 0.096±0.008 |
| 1, 3 | 0.905±0.028 | 0.066±0.010 |
| 2, 3 | 0.963±0.017 | 0.116±0.010 |

Correlations between breeding values for non-EA rate from data set 5 and the Israeli breeding index, PD16, and the other economic traits computed for Israeli Holsteins are given in Table 5. All correlations were significantly different from zero, except for the correlations with fat and protein production. The correlation with PD16 was 0.115. Thus selection for the index should have resulted in a decrease in abortion rate. The regression of the breeding value for non-EA rate on the cows’ birth year was 0.083% per year (P<0.001), that is a decrease of close to 0.1% per year. The highest correlation was with female fertility, 0.75; but the correlation with herd-life was also 0.3. Correlations among breeding values tend to slightly underestimate the actual genetic correlations, due to incomplete reliabilities of the evaluations.

**Table 5.** Correlations between breeding values for non-abortion rate from data set 5 and the Israeli breeding index (PD16) and the other economic traits computed for Israeli Holsteins.

| Traits | % of index | Number of bulls | Correlation |
| --- | --- | --- | --- |
| PD16 | 100 | 1693 | 0.115*** |
| Milk (kg) | 0.00 | 1693 | -0.077** |
| Fat (kg) | 21.20 | 1693 | -0.026 |
| Protein (kg) | 37.32 | 1693 | -0.044 |
| SCS<sup>1</sup> | 10.98 | 1693 | -0.196*** |
| Female fertility (%) | 14.38 | 1693 | 0.749*** |
| Herd-life (days) | 9.58 | 1693 | 0.296*** |
| Persistency (%) | 4.24 | 1693 | 0.078** |
| Dystocia, maternal (%)<sup>a</sup> | 1.27 | 1693 | -0.179*** |
| Stillbirth, maternal (%)<sup>a</sup> | 1.03 | 1693 | -0.246*** |
| Dystocia, direct (%)<sup>1</sup> | 0.00 | 1592 | -0.105*** |
| Stillbirth, direct (%)<sup>1</sup> | 0.00 | 1592 | -0.112*** |

The relative contribution of each to the index and the numbers of bulls with evaluations with reliabilities > 0.5 for each trait are also listed.
<sup>a</sup> Negative values are economically favorable.
\* significant, $p < 0.05$ ; \*\*, significant, $p < 0.01$ , \*\*\*, significant, $p < 0.001$ .

Mean phenotypic and breeding values of non-EA rate for first parity cows by birth year are given in Fig 3. Although the overall regression of breeding value was positive for non-EA rate, EA rate increased until 1994, and then decreased beginning in 2002. These changes correspond to changes in the Israeli breeding index. Until 1996 the index included only milk production traits. Somatic cell score was added in 1996, and female fertility in 2000; which now accounts for 14% of the index. A lag of ∼2 years between inclusion of a trait in the index and a change in the effective direction of selection is expected. The regression of breeding value on birth date for cows born since 2002 was 0.53% per year, as opposed to 0.083% since 1983. No clear trend is evident for the phenotypic means of first parity EA rate.

**Fig 3.**
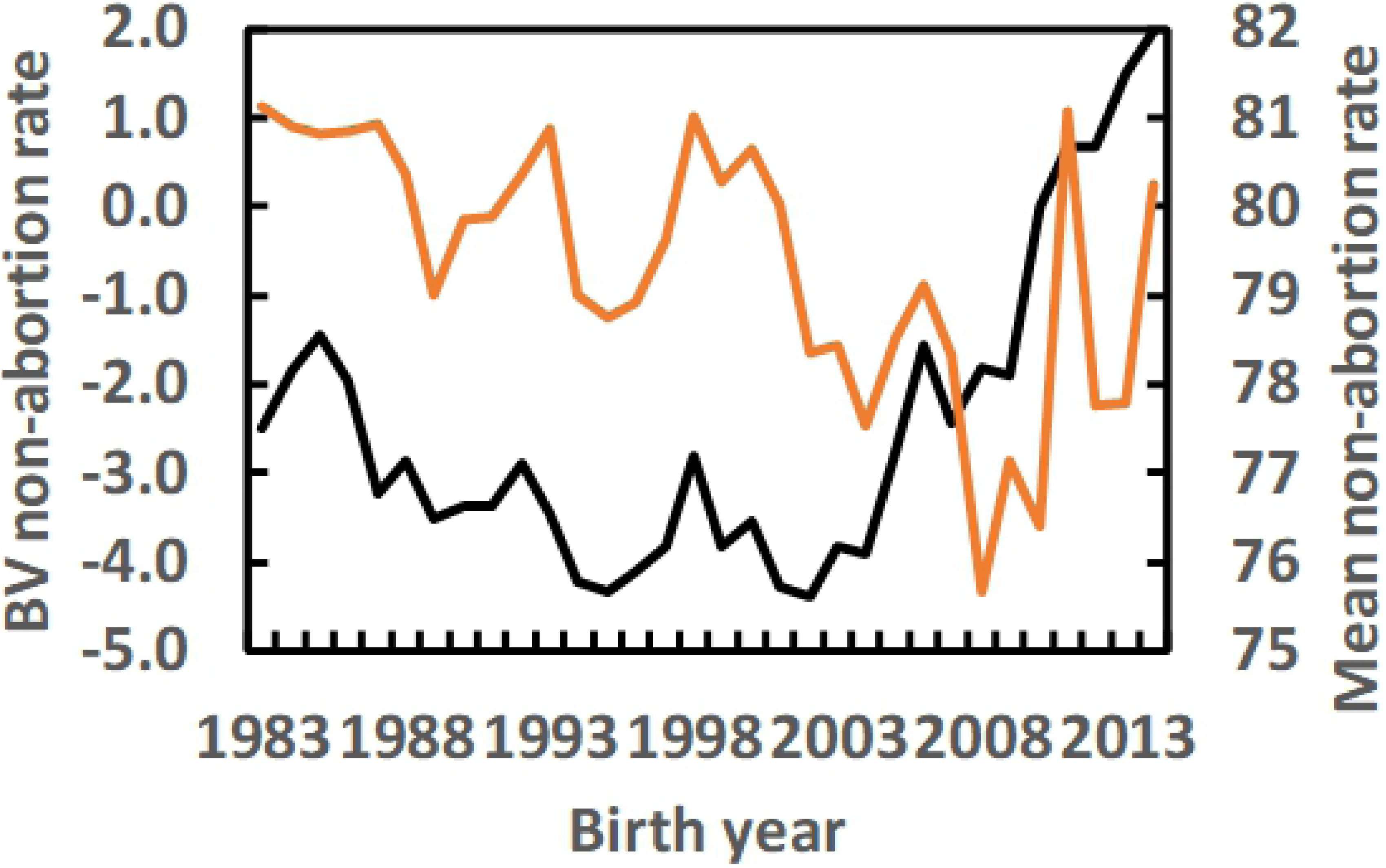
Mean breeding values of non-abortion rate and mean non-abortion rate of first parity cows by birth year. 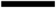, Mean breeding value for non-abortion rate; 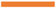, Mean non-abortion rate. A score of 0 = abortion, and 100 = normal pregnancy. Genetic evaluations are computed relative to cows with born in 2010.

The “Manhattan Plot” for the genome wide association study (GWAS) results for EA rate are given in Fig 4, and the markers with the lowest probability values are presented in Table 6. There were eight markers with nominal probabilities < 10^-11^. All of these markers has probabilities <10^-6^ after permutation analysis and correction for multiple testing. Of the 8 markers listed, the 2 markers on chromosome 7 and the 2 markers on chromosome 17 are clearly due to a single quantitative trait locus segregating on each chromosome, since the distance between the two markers on each chromosome is <100,000 base pairs. Each of these 8 markers explained between 5 and 4% of the variance for the genetic evaluations for EA rate, and > 2.5% of the variance for CR. None of these chromosomal regions were found to have significant effects on cow conception rate or daughter pregnancy rate in the analysis of the US Holstein population by the posteriori granddaughter design [30].

**Table 6.** Markers with the lowest probability values for early abortion rate.

| Location |  | SNP | Early abortion rate |  |  | Conception rate |  |  |
| --- | --- | --- | --- | --- | --- | --- | --- | --- |
| Chr. | Base pairs |  | Beta<sup>a</sup> | R<sup>2b</sup> | P<sup>c</sup> | Beta | R<sup>2</sup> | P |
| 19 | 61503930 | Hapmap43271-BTA-46356 | 1.56 | 0.051 | 4.96E-15 | 0.89 | 0.058 | 1.30E-18 |
| 8 | 24608595 | Hapmap41408-BTA-103152 | 1.52 | 0.047 | 6.08E-14 | 0.83 | 0.048 | 1.34E-15 |
| 17 | 26712567 | BTA-46662-no-rs | -1.95 | 0.047 | 6.60E-14 | -1.02 | 0.045 | 1.19E-14 |
| 21 | 46984914 | BTA-52458-no-rs | 1.49 | 0.047 | 6.63E-14 | 0.60 | 0.026 | 4.51E-09 |
| 17 | 26735031 | Hapmap41875-BTA-46663 | -2.23 | 0.045 | 1.81E-13 | -1.26 | 0.050 | 3.43E-16 |
| 24 | 53790841 | BTA-58638-no-rs | 1.77 | 0.043 | 4.99E-13 | 0.84 | 0.035 | 1.47E-11 |
| 7 | 22920391 | BTB-01966013 | 1.49 | 0.042 | 1.01E-12 | 0.68 | 0.031 | 2.07E-10 |
| 7 | 22996615 | BTB-01398686 | 1.47 | 0.041 | 1.95E-12 | 0.66 | 0.029 | 6.26E-10 |

Markers are sorted in descending order of the probability to reject the null hypothesis of no effect on early abortion rate. The substitution effects and coefficients of determination are given for each marker for early abortion and conception rate.
<sup>a</sup>The allele substitution effects in transmitting value trait units.
<sup>b</sup>Coefficient of determination.
<sup>c</sup>The nominal probability for the hypothesis of no effect. All genome-wide probabilities were $<10^{-6}$ .

**Fig 4.**
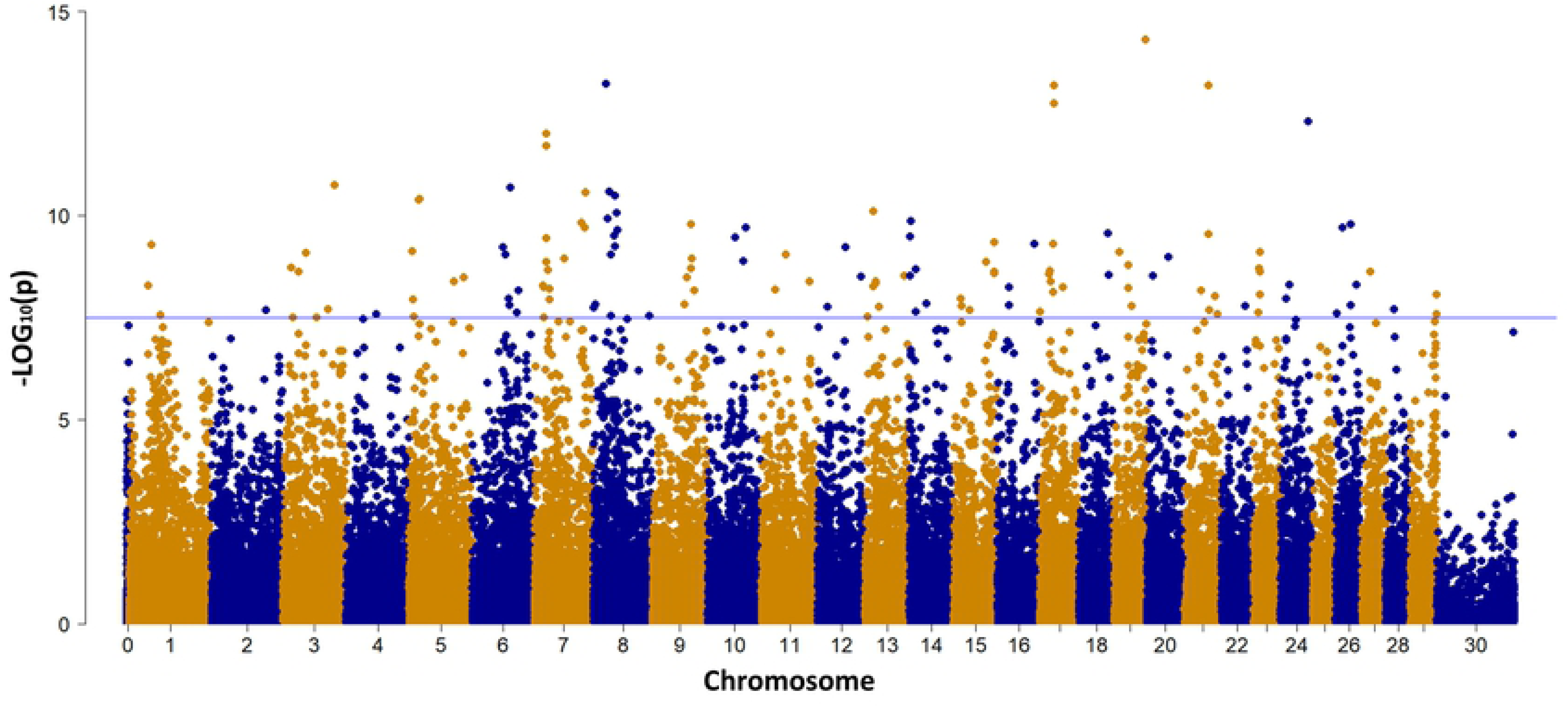
GWAS Manhattan plot for EA rate. Dots represent each marker. Chromosomal positions are on the X-axis, and nominal –log10 P-values are on the Y-axis. The blue line denotes the genome-wide significance threshold of 0.05, as derived from one million data permutations and correction for multiple testing.

While all of the 30 markers with probabilities < 10^-8^ for EA are also significantly associated with CR, only subset of the markers with probabilities < 10^-12^ for CR are also significantly associated with EA (Fig 5). The effects of these 30 markers on EA on their effects on CR is plotted in Fig 6. The regression was 2.0 with a coefficient of determination of 0.97. Thus the substitution effect for EA was generally twice the effect for CR.

**Fig 5.**
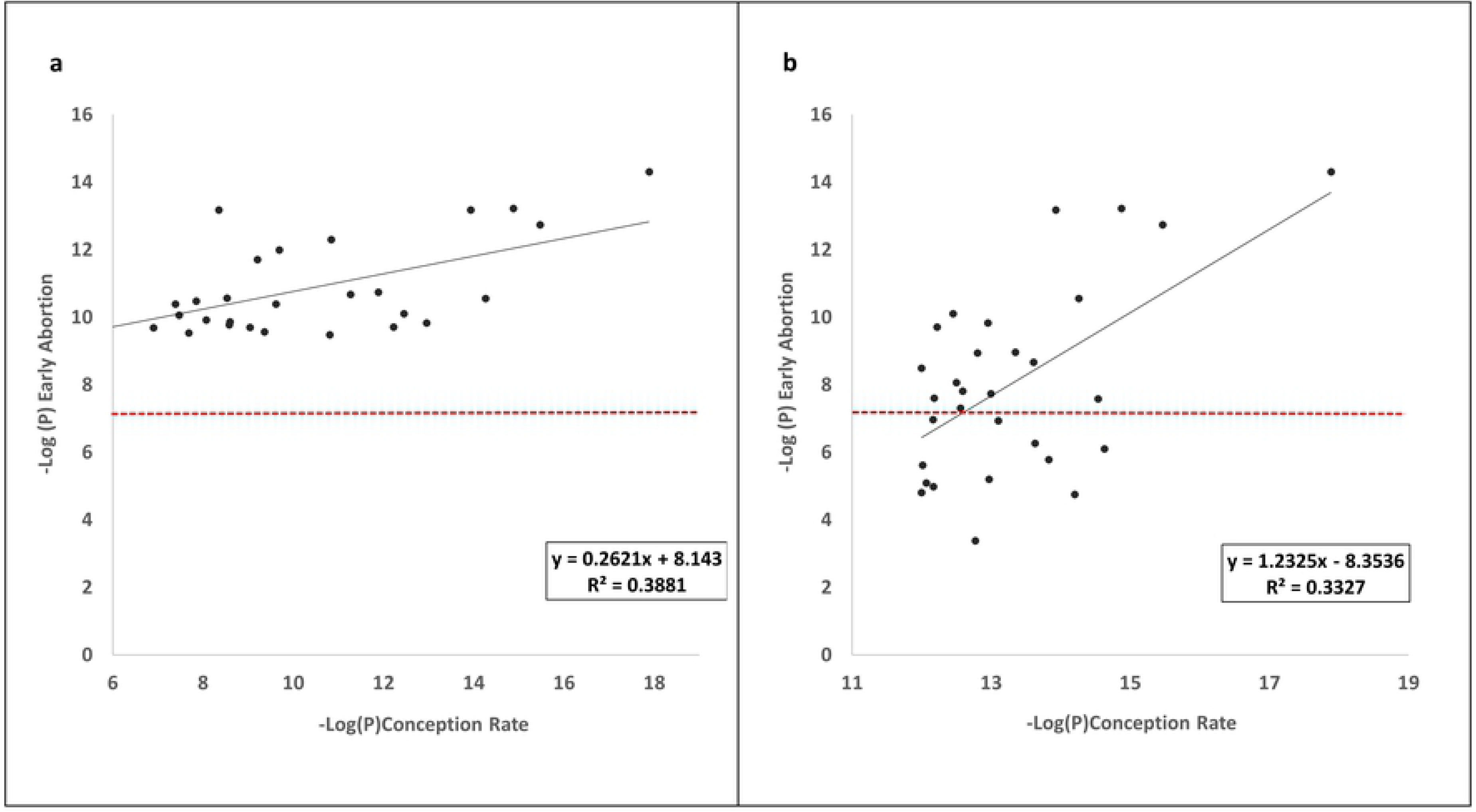
Comparison between the GWAS results for early abortion rate and conception rate. a) The 30 markers with the lowest nominal probabilities for early abortion rate as a function of the corresponding probabilities of these markers for conception rate. b) The 30 markers with the lowest nominal probabilities for conception rate as a function of the corresponding probabilities of these markers for early abortion rate. The red dashed line denotes the genome-wide significance threshold of 0.05 for early abortion as derived from one million data permutations and after correction for multiple testing.

**Fig 6.**
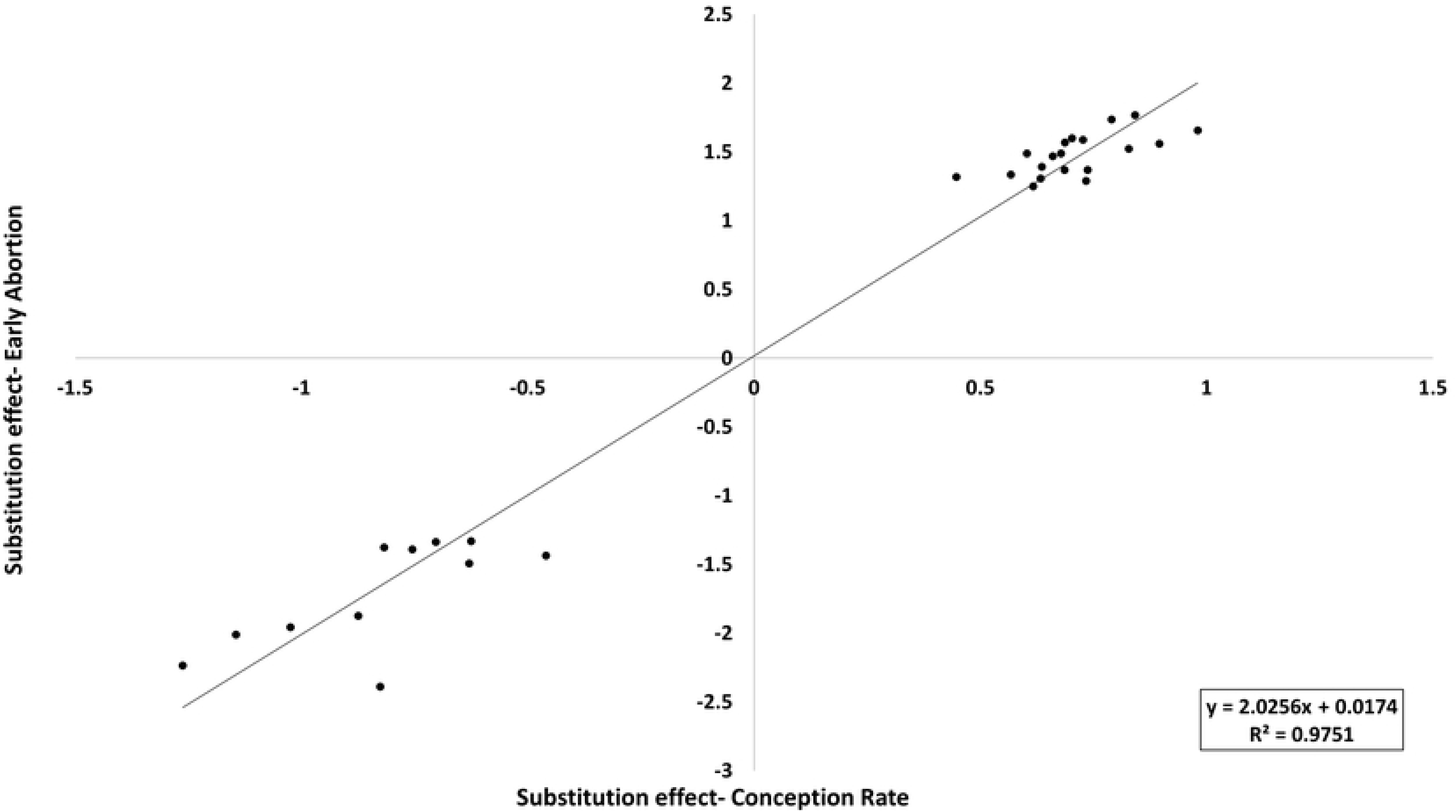
The substitution effects for early abortion rate of the 30 markers with the lowest nominal probabilities as a function of the substation effects for the same markers for conception rate.

## Discussion

Reduce fertility is a major concern in humans, and a highly important economic trait in dairy cattle. Previous studies suggested that a large portion of recorded non-conceptions in human and cattle are apparently the result of unrecognized EA [8, 9, 21]. In this study we used the extensive records of the Israeli dairy cattle population to study the genetics and the genomics of EA and to assess its association with CR.

CR can be affected by the service sire and/or by the dam inseminated. In cattle, the service sire can effect CR either via the quality of semen, or via genes that reduce embryo survival rate. With respect to EA, the effect of the sire is limited to genes affection embryo survival rate. The variance of the service sire effect on CR is approximately three-fold the variance of the service sire effect on EA rate (Table 3). This suggests that most of the service sire effect on CR is due to semen quality, otherwise the service sire variance component should have been of similar magnitude to the variance component for EA. Although several recessive lethal genes have been detected that cause EA [25], their effects on the genetic variance of EA rate appears to be minimal. This is not surprising, considering that the lethal allele is always quite rare. Generation of homozygotes for rare alleles is generally due to inbreeding, but inbreeding is carefully monitored in the Israeli dairy cattle population. The inbreeding from each potential mating is checked by the inseminator, and matings that result in >3.125% inbreeding are generally rejected [31].

Only 2 previous studies attempted to estimate heritability of EA in dairy cattle, and both are somewhat problematic. In Bamber et al. [32] pregnancy loss was determined by an initial pregnancy diagnosis 26 to 33 d after AI followed by determination of loss of that embryo at a subsequent diagnosis 14 to 39 d later, but only 3,775 cows were included in the study. Due to the relatively small sample, confidence intervals for the genetic parameters were so wide, as to render the results virtually meaningless. They found a heritability of 17%, which was not significantly different from the value of ∼10% in the current study. However, they found that the service sire variance was 16% of the total variance. This is clearly at variance with the current study, and was also considered difficult to explain by [32]. In Carthy et al. [33] EA was assumed to have occurred if pregnancy was determined by ultrasound examination, and the embryo was later deemed to be unviable by a later examination. However, ultrasound examinations were performed at various time points postpartum at the discretion of the producer. On a sample of 43,473 lactations they found heritability of only 2%, but repeatability of 66%. Part of the discrepancy to the current study may be due to the fact that the time periods for determination of both pregnancy and abortion were not consistent across records. Mean embryo loss was only 8%, as compared to 22% on first parity in the current study.

The genetic trend for EA rate corresponds to changes in the Israeli breeding index, as noted previously. However, the phenotypic trend is not similar to the genetic trend. Several factors can possibly explain this, including climatic variation, and the fact that the phenotypic means were computed only for first parity cows.

The GWAS results show that the markers with the lowest probability values for EA are also included among the markers with the lowest probability values for CR, but not vice versa (Fig 5). In addition, the regression of the substitution effects of the significant EA markers on their substitution effects for CR is ∼2 (Fig 6). This suggests that the genetic factors that affect EA are likely a subgroup of factors affecting the CR. The markers that explain the most variance for EA suggest possible new insights on the polygenic architecture of this trait. For instance, by investigating the genomic area flanking these markers (Table 6), we found that the marker explaining the most variance (Table 6, Fig 7a) is located within the ABCA9 gene, that is found within an ABC genes cluster (Fig 7 a). The ATP-binding cassette (ABC) family are the major class of primary active transporters in the placenta. ABC proteins are reported to be important in efflux of xenobiotics and endogenous substrates like lipids, sterols and nucleotides. Recent studies provided evidence that ABC genes protect placental tissue by preventing accumulation of cytotoxic compounds, which is important in complicated pregnancies, such as in inflammatory or oxidative stress [34]. Thus, our finding of genetic variation within ABC genes cluster suggest that differences in EA predisposition might involve different sensitivity for oxidative stresses during the first trimester, mediated by ABC genes.

**Fig 7.**
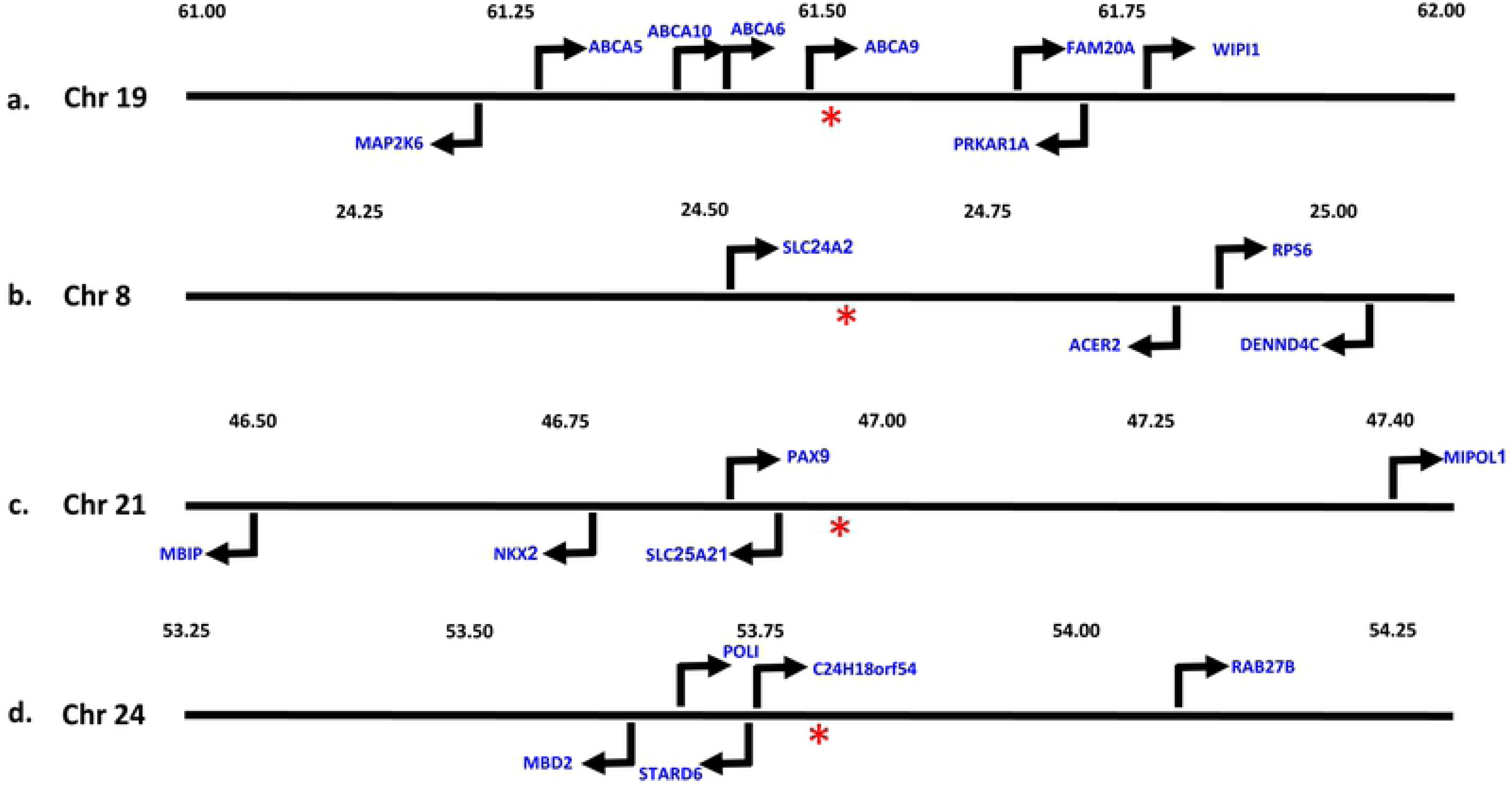
Schematic representation of the genomic area flanking four of the markers with the lowest nominal probability values (Table 6). The arrows represent the genes position and the strand orientation. The red asterisk represents the marker positions for a) Hapmap43271-BTA-46356 located on chromosome 19 b) Hapmap41408-BTA-103152 located on chromosome 8 c) BTA-52458-no-rs located on chromosome 21 and d) BTA-58638-no-rs located on chromosome 21.

The significant marker on chromosome 21 is adjacent to the genes PAX9 and NKX2-1 (Fig 7c). Previous works suggested that PAX9 is required for the chondrogenic differentiation of sclerotomal cells during embryogenesis [35, 36], and that NKX2 is required for the embryonic development of cholinergic septohippocampal projection neurons [37]. Moreover, according to PathCards, NKX2-1 is part of the embryo pre-implementation path [38]. The fact that this marker is associated with a major effect on EA suggests that altered predisposition for EA might be mediated by genes that regulating embryo development.

Since fertility is a major economical trait in dairy cattle [18], our results suggest that EA, as defined in this study, should be considered for inclusion in the commercial selection index. Shook [39] listed the criteria that a potential trait must meet in order to be included in the selection objective. First, it should have an economic value. Second, the trait must have sufficiently large genetic variation in relation to its economic value and heritability. Third, the trait should be measurable at a low cost and consistently recorded. An indicator trait may be favored if it has a high genetic correlation with the economically important trait, is easier to record, has a higher heritability than the economic trait, or can be measured earlier in life [40]. The classic example of selection on an indicator trait in dairy cattle is somatic cell score as an indicator trait for mastitis [39]. The two traits have a relatively high genetic correlation, but somatic cell score has heritability of 10-20%, as opposed to 2-6% for clinical mastitis [39, 41]. Selection on EA rate, as defined in the current analysis is even more attractive as an indicator trait for female fertility; in that there is no requirement to generate new data. Considering the apparent high genetic correlation, and the fact that the heritability for EA rate is three-fold the heritability for most measures of fertility, genetic progress for fertility will be higher via selection for EA rate.

## Materials and methods

### Data sets analyzed

Eight data sets were analyzed. A basic description of the data sets is given in Table 1, and the numbers of animals and levels of effects included in each data set are given in Table 2. The first data set included conception status of first parity cows Israeli-Holstein cows with calving dates from Jan. 1, 2007, through Dec. 31, 2016, and at least one insemination. Fertility data in Israel is unique in that cows that are not re-inseminated within 60 days are checked for pregnancy by a veterinarian [19]. The following records were deleted:

**Table 1.** Basic description of the seven data sets analyzed.

| Data Set | Description | Analysis Type<sup>a</sup> | Parities | Freshening years included |  |
| --- | --- | --- | --- | --- | --- |
|  |  |  |  | Beginning | End |
| 1 | Conception rate, all cows | VC | 1 | 2007 | 2016 |
| 2 | Non-abortion rate | VC | 1 | 2007 | 2016 |
| 3 | Conception rate without abortions | VC | 1 | 2007 | 2016 |
| 4 | Non-abortion rate | VC | 1-3 | 2007 | 2016 |
| 5 | Non-abortion rate | SAM | 1-3 | 1985 | 2016 |
| 6 | Female fertility | MAM | 1-5 | 1985 | 2018 |
| 7 | Non-abortion rate, GWAS | GWAS | 1-3 | 1985 | 2016 |
| 8 | Female fertility, GWAS | GWAS | 1-3 | 1985 | 2016 |

<sup>a</sup> VC = variance component analysis, SAM = single trait animal model, MAM = multi-trait animal model, GWAS = genome-wide association study.

**Table 2.** Number of animals with records, ancestors and levels of effects in the analysis models. Data sets are defined in Table 1.

| Data Set | Number of levels or animals |  |  |  |  |  |
| --- | --- | --- | --- | --- | --- | --- |
|  | Insemination month | Service sire | Herd-year-seasons | Genetic groups | Animals with records | Ancestors |
| 1 | 12 | 569 | 13699 | 2 | 260782 | 142297 |
| 2 | 12 | 554 | 12802 | 2 | 118039 | 123600 |
| 3 | 12 | 568 | 13552 | 2 | 233037 | 142816 |
| 4 | - | - | 13551 | 2 | 182758 | 136187 |
|  | 12 | 554 | - | - | 118039 | - |
|  | 12 | 542 | - | - | 77193 | - |
|  | 12 | 431 | - | - | 51234 | - |
| 5 | - | - | 48757 | 84 | 571988 | 181478 |
| 6 | - | - | 50536 | 80 | 861939 | 143649 |
| 7 | - | - | - | - | 1179 | - |
| 8 | - | - | - | - | 1297 | - |

Data sets are defined in Table 1.

1. Cows that were daughters of foreign bulls.
2. Cows with first insemination by foreign bulls.
3. Cows with first insemination ≤30 and ≥135 days after parturition.
4. Cows for which pregnancy could not be determined for the first insemination. This chiefly included cows that were not re-inseminated, and were culled prior to determination of pregnancy by veterinary inspection.

The second data set was a subset of the first set, and included all cows that were either recorded as pregnant on the first insemination, and with pregnancy length ≤ 290 days; or cows that were recorded as “open” on the first insemination, and re-inseminated between 49 and 100 days after the first insemination. The latter group was assumed to represents cows with EA.

The third data set included all cows in the first data set recorded as pregnant, and cows that were recorded as open and re-inseminated prior to 49 days after the first insemination. This data set was assumed to represent cows that either became pregnant or remained open after the first insemination. Although the objective was to eliminate cows with EA, the latter group of cows clearly include a small fraction of cows that aborted, and still displayed estrus prior to 49 days after the first insemination.

Cows were included in the fourth data based on the same criteria as the second data set, but cows in second and third parities were also included. Since cows with second inseminations < 49 days or >100 days after first insemination were deleted for each parity, some cows had records on one, two or three parities, and for any specific parity a record could be included or deleted. Variance components were computed for all four data sets for a binary trait. The analyses included all parents and grandparents of cows with valid records.

The fifth data set was similar to the fourth data set, except that it included all first through third parity cows with freshening dates from Jan. 1, 1985 through Dec. 31, 2016, which either became pregnant on first insemination or were re-inseminated between 49 and 100 days after the first insemination. This data set also included cows that were daughters of foreign bulls, and even bulls of breeds other than Holsteins, although these cows were only about 1% of all the cows. This data set was analyzed by the individual animal model in order to compute genetic evaluations for all bulls with genotypes for medium or high density SNP-chips. As in the previous data sets, the animal model analysis included all parents and grandparents of cows with valid records.

The sixth data set included all valid first through fifth parity records for female fertility with freshening dated from Jan 1 1985 through May 31, 2018. All cows that were inseminated at least once were included. As described previously [42], fertility was scored as the inverse of the number of inseminations to conception in percent. For cows that were culled prior to conception, the expect number of inseminations to conception was computed. As in data set 5, cows that were daughters of foreign Holstein bulls and bulls from breeds other than Holsteins were included, and the data set included all parent and grandparents of cows with valid records. This data set was analyzed by the multi-trait animal model, with each parity considered a separate trait, as described by [43]. The separate parity evaluations were combined into a multi-parity index, based on the economic value of each parity.

The seventh data set included all bulls with genetic evaluations from the analysis of the fifth data set with reliabilities > 50% and genotypes for one of the medium or high density SNP-chips. This data set was used to compute the GWAS analysis for frequency of EA. The final data set included all bulls with genetic evaluations from the sixth data set with reliabilities > 50% and genotypes. A GWAS analyses was also computed on this data set, and compared to the GWAS for EA rate.

### Statistical analyses

Variance components were estimated for data sets one through four using the AIREMLf90 program [44]. The trait analyzed in data sets one and three was CR for the first insemination, and for data sets two and four the trait analyzed was EA rate, under the assumption that re-insemination between 49 and 100 days indicates an early abortion. Both traits were scored dichotomously, with non-conception or abortion scored as zero, and pregnancy scored as 100. A single trait animal model was assumed for data sets one through three, and a multi-trait animal model was assumed for data set four. For data set four, the three parities were considered three separate traits. In addition to the random additive genetic effect of the cow calving, and service sire for the first insemination, all models included the effect of insemination month and herd-year-season as fixed effects. Two seasons were defined for each herd-year beginning in April and October of each year. For data sets one through four, two genetic groups were defined for animals with unlisted parents, one for males and one for females.

Heritability was defined as the ratio of the additive genetic variance to the sum of additive genetic, service sire and the random residual variances. Genetic and environmental correlations among the parities were computed for data set four. Genetic correlations were the correlations among the additive genetic effects, and the environmental correlations were the correlations among the residual effects. The AIREMLf90 program also computes solutions for all effects included in the analysis model and standard errors for all variance components and the heritabilities and the correlations.

Data set five was analyzed by a single trait animal model with all three parities considered the same trait, as described by [42]. Thus the model included a random permanent environmental effect in addition to the additive genetic effect. These effects differ in their variance structure in that only the additive genetic effect included the relationship matrix. The assumed ratios of variance between the residual and the additive genetic and permanent environmental effects were both 9. Variances of abortion rate were lower in first and second parities, due to lower abortion rates. Therefore, first and second parity records were each multiplied by a factor greater than unity to obtain equal phenotypic variances for all three parities. The adjusted records were then adjusted for the mean effects of parity and insemination month by subtracting the means of the parity-insemination month classes from each record. In addition to the additive genetic and the permanent environmental effects, the model included the herd-year-season effect, as described previously; a parity-by herd type effect and a genetic group effect. Two herd-types were defined; “Moshav” (family farms) and “Kibbutz” (communal herds). Although the records were pre-adjusted for parity effects, a residual effect could remain after accounting for all the effects included in the model.

In the analysis of data set 5, 84 groups were defined based on the sex of the animal with unknown parents, which parent was unknown, and the birth year. In addition, separate groups were defined for sire of cows of breeds other than Holstein. Although only a very small fraction of the cows was sired by bulls of other breeds, these bulls were a significant fraction of the total number of bulls, and an even larger fraction of the bulls with unknown parents.

The overall genetic trend was computed as the regression of the cows’ breeding values on their birth dates, for all cows born since Jan. 1, 1983. Yearly means of first parity non-abortion rate and the breeding values of cows by birth year, relative to cows born in 2010, were computed. Reliabilities of the breeding values of all animals included in the analysis were estimated by the method of [45]. There were 1701 sires with reliabilities > 0.5 for EA rate. Correlations were computed between the breeding values for EA rate for these sires and the current Israel breeding index, PD16 and 11 economic traits analyzed in Israel. Breeding values for these traits other than female fertility were computed as described previously [43, 46-48].

### Genome-wide association studies

A total of 1749 Israeli Holstein bulls were genotyped. Since genotyping of these sires were performed using several SNP-chip platforms, we filtered-in only those markers that were covered in more than 90% of the tested cohort. Approximately 41,000 SNPs were retained. Genome-wide associations were computed for the sires’ transmitting abilities (½ of the breeding value) for EA rate and CR (Table 1 and 2, data sets 6 and 7). Of the genotyped bulls, there were 1179 and 1297 with genetic evaluations for EA rate and CR, with reliabilities > 50%, respectively. The additive substitution effects, the coefficients of determination (R^2^) and the nominal probabilities for the hypothesis of no effect were computed by using plink software [49]. Genome-wide probabilities were estimated by generating one million permutations of genotype data against the genetic evaluations. Thus, the minimal genome-wide probability was <10^-6^ if the substitution effect obtained from the actual data was greater than all of the permutation effects.

## Acknowledgments

This research was supported by grant number 58-8042-5-063F from the U.S.-Israel Binational Agricultural Research and Development (BARD) Fund, and by a grant from the Israel Dairy Board. We thank Ignacy Misztal and Shogo Tsuruta for use of the AIREMLF90 program, and Michael van Straten for useful discussions.

